# Isolating Poroelastic and Viscoelastic Mechanisms of Soft Tissues and Hydrogels Through Sequential Microscale Indentation Testing: New Applications of Indentation Theory for Microscale Characterization

**DOI:** 10.1101/2024.06.16.599204

**Authors:** Muhtadi Munawar Zahin, Abeer Al Barghouthi, Darryl A. Dickerson

## Abstract

Soft hydrated materials, including biological tissues and hydrogels, exhibit complex time-dependent mechanical behaviors due to their poroelastic and viscoelastic properties. These properties often manifest on overlapping time scales, making it challenging to isolate the individual contributions of poroelasticity and viscoelasticity to the overall mechanical response. This study presents a novel semi-analytical model for characterizing these properties through sequential microscale load relaxation indentation testing. By extending existing theories, we developed a poroviscoelastic framework that enables the deconvolution of poroelastic and viscoelastic effects. Using this model to fit sequential microscale indentation data, we characterized porcine heart and liver tissues, as well as collagen and GelMA hydrogels, revealing distinct differences in their poroelastic and viscoelastic parameters. Our findings demonstrate that this approach not only provides rapid and detailed insights into the mechanical properties at the microscale but also offers significant advantages over traditional methods in terms of speed, computational efficiency, and practicality. This methodology has broad implications for advancing the understanding of tissue mechanics and the design of biomimetic materials for tissue engineering applications.

**Statement of Significance:** This study introduces a novel approach to understanding the mechanical behavior of soft hydrated materials, like tissues and hydrogels. This study introduces a semi-analytical model to describe the time dependent behavior and a practical approach to distinguish between poroelasticity and viscoelasticity at the microscale. By providing this model along with a rapid and efficient characterization method, our approach enhances understanding of time-dependent mechanical behaviors critical for soft tissue mechanics and biomaterials design.

## 1 Introduction

Soft tissues and extracellular matrix mimics (e.g., hydrogels), which in this work we will refer to collectively as soft hydrated materials, exhibit time dependent behavior under mechanical loading. This time dependent behavior is a ubiquitous characteristic of soft hydrated materials that has implications both for how we understand soft tissues and for how we approach tissue engineering, particularly given the common use of hydrogels as biomaterial components for soft tissue engineering. In soft tissues, time dependent mechanical properties provide insight into the health status and age of tissues. For example, while the change in elastic response of tissues is the most widely studied biophysical marker of cancer [1–3], substantially different time dependent responses are observed in cancerous tissues in comparison to normal tissues [4,5], which may serve as a better marker for health state of the tissue [5]. Differences in time dependent behaviors are also observed in entheses, soft to hard tissue interfaces, in healthy and osteoarthritic joints [6,7]. Age-related and damage-related changes in time-dependent properties have been observed in multiple tissues including skin, tendons, cartilage, brain, and muscle [7– 19]. In developing models for tissue behavior and materials for tissue regeneration, time dependent mechanical properties have been found to be an important driver of a number of cellular responses. For example, differences in relaxation response within hydrogels have been demonstrated to influence mesenchymal stem cell fate [20,21]. Similarly, these properties have been shown to influence the organization of epithelial tissues and organization of pluripotent stem cell clusters in hydrogels [22,23]. It is evident that increasing our understanding of time dependent mechanical properties provides important information about the baseline state of soft tissues and can provide new insights into how we can harness these biophysical cues for tissue engineering efforts.

The time dependent properties of soft hydrated materials arise from two distinct fundamental mechanisms related to their composition: poroelasticity and viscoelasticity. Soft hydrated materials have a multicomponent nature – they are porous, heterogenous solid polymeric networks which are filled with large volumes of interstitial fluid. The time dependent behavior observed in soft hydrated materials under loading can be attributed to the resultant effect of the different ways in which the solid and fluid components are rearranged as a result of that loading. The first mode of rearrangement occurs due to response of interstitial fluid within soft hydrated materials. Loading leads to pressurization of the fluid within the pores followed by rearrangement of the interstitial fluid relative to the pores. Under a constant displacement, the fluid moves until the pressure releases to an equilibrium state. This mode of relative rearrangement of the interstitial fluid and the porous solid skeleton defines poroelasticity. Poroelasticity affects solute movement at multiple scales, thus influencing nutrient transport to and from cells, as well as local signaling between cells. Additionally, fluid movement provides other biophysical cues including hydrostatic pressures and shear stresses acting on cells [24,25]. The solid skeleton of soft hydrated materials can undergo a second mode of rearrangement, characterized by the sliding of solid components, polymeric unfolding, and the relatively rapid dissociation of weak bonds. These processes contribute to the dissipation of some of the energy of deformation, resulting in the intrinsic viscoelastic response [26–28]. Cells deform and exert forces on the matrices in which they reside, experiencing the energy dissipation due to viscoelasticity, which influences cell spreading behavior [29,30], mechanosensitive nuclear localization of transcription regulators [30–33], and cellular differentiation outcomes [20].

Both poroelastic and viscoelastic behaviors co-exist within soft hydrated materials at the tissue-scale and at the cell length scale. The hierarchical structure of tissues means that mechanical behaviors may vary across different length scales. Thus, tissue scale examination of mechanical properties may not be sufficient. Examining how these behaviors manifest at the cell scale is critical to provide a baseline understanding of soft hydrated materials and to inform design of biomimetic materials for both in vitro models and eventual therapeutic use, thus motivating the use of nanoindentation [34,35]. However, the two mechanisms of time-dependent behavior may occur on the same time scale, making it challenging to distinguish these distinct mechanical properties. In the current corpus of literature, there have been several approaches to meet this challenge. One approach is to use inverse parameter identification with finite element software. Such methods, however, are currently computationally expensive and may be subject to the intricacies of the individual finite element software. Islam and Oyen examined how different models of time-dependent behavior fit cell-scale relaxation responses to classify the behavior as having more poroelastic or viscoelastic character [34]. Other researchers have used a similar fitting approach to extract time points where the dominant time-dependent behavior changes [36]. While useful, these approaches do not permit the true isolation of the contributions of poroelasticity and viscoelasticity in soft hydrated materials.

The aim of the present study is to formulate a new theoretical foundation for poroviscoelastic characterization and to present a practical sequential indentation method to isolate the poroelastic and viscoelastic properties of soft hydrated materials. In this work, we extend Wang et al’s recently developed poroelastic large relaxation indentation theory [37] to a porovisceolastic theory that accounts for ramp indentation within the theoretical formulation in contrast to many studies where step-loading is assumed for analysis but is possible to practically implement. We then use this theoretical foundation along with scaling arguments to provide a new framework for characterization of soft hydrated materials, deconvolving poroelasticity and viscoelasticity and identifying the individual contributions of each to time dependent behavior. Using this framework, we characterize several soft hydrated materials (porcine heart, porcine liver, gelatin methacryloyl hydrogels, and collagen hydrogels) to illustrate the utility of this new approach.

## 2 Materials and Methods

### 2.1 Theory

#### 2.1.1 Foundational Poroelastic Indentation Theory for Extended Load Relaxation Ranges

In order to describe the time dependent mechanical behavior of soft hydrated materials, we build on Wang and colleagues’ improved indentation theory that accounts for a larger range of load relaxation magnitudes that may be observed in soft hydrated materials. This theory extends Biot’s poroelastic theory and classical indentation theory and is described here in brief. We begin by considering the soft hydrated material as a poroelastic half-space being indented by a sphere. We assume the sphere to have a much greater modulus than that of the soft hydrated material and can therefore be considered a rigid body. Contact between the sphere and the material surface is considered to be frictionless. Assuming the soft hydrated material consists of a porous linearly elastic solid saturated with a fluid phase and modeling contact using the Hertz solution, Wang defined the indentation force as a function of time during an indentation ramp and hold as follows:

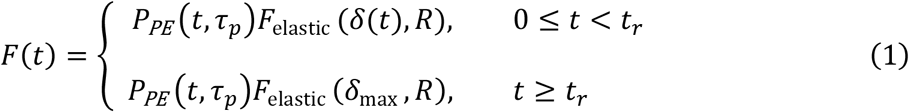

where *δ*(*t*) is the indentation depth as a function of time, *R* is the radius of the indenter, *F*_elastic_ (*δ*(*t*), *R*) is the elastic force due to indentation, *t*_*r*_ is the ramp time, *P*_*PE*_ (*t, τ*_*p*_) is the poroelastic pressurization function, and *τ*_*p*_ is the poroelastic time constant. We define the elastic force here using Hertz contact theory with a spherical indenter as:

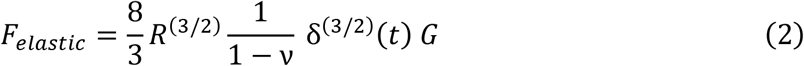

Where *G* is the equilibrium shear modulus of the solid matrix and *v* is the drained Poisson’s ratio. Wang and colleagues define the poroelastic pressurization function as:

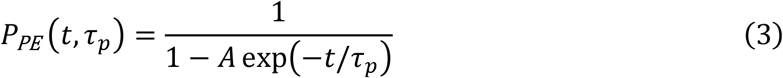

Where A is a material constant. The poroelastic time constant *τ*_*p*_ can be written as:

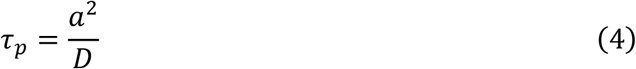

Where D is the diffusivity and a is the radius of contact which is calculated as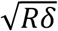. We interpret the Wang formulation as describing poroelastic load as pressurization due to fluid constriction within the solid matrix which is quantified by a multiplier that reaches a theoretical constant value 1/(1 − *A*) under instantaneous step loading. This formulation captures how that pressurization constant changes under more realistic ramp loading through the exponential term in the denominator. We recast the pressurization function into a more general form as follows:

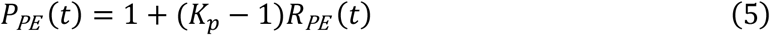

Where *K*_*p*_ is the pressurization constant equation to 1/(1 − *A*) and *R*_*PE*_ (*t*) is the poroelastic relaxation function. *P*_*PE*_ (*t*) represents the load magnification over the elastic force predicted by Hertz contact theory which also encompasses fluid diffusion during the load ramp. There are two important observations from this formulation. First, as the ramp time increases, the load magnification decreases as fluid diffuses away during the ramp. Second, at longer time periods, the equilibrium response of this formulation is equivalent to the elastic indentation response.

The time-dependent portion of models of poroelastic relaxation typically have some combination of exponential terms that take the following form:

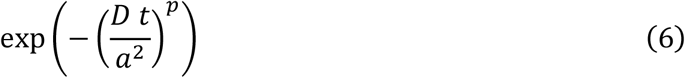

where *t* is time, *D* is diffusivity, *a* is the contact radius, and *p* is a positive constant that determines the specific form of time dependence. Thus, we can generally describe poroelastic relaxation during the hold as follows:

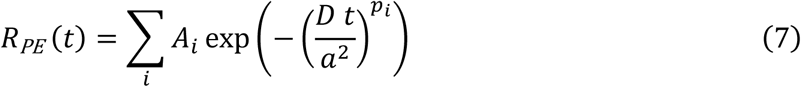

Where the sum of the *A*_*i*_ = 1 for the prior equation. For the ramp phase of loading, the radius of contact *a* is a function of time. We will assume the application of a linear indentation ramp of constant velocity *v*_*r*_ = *δ*_max_ /*t*_*r*_ and we define *δ*(*t*) = *v*_*r*_ *t*.

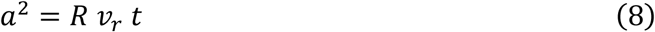

Substituting into the poroelastic relaxation equation, we get:

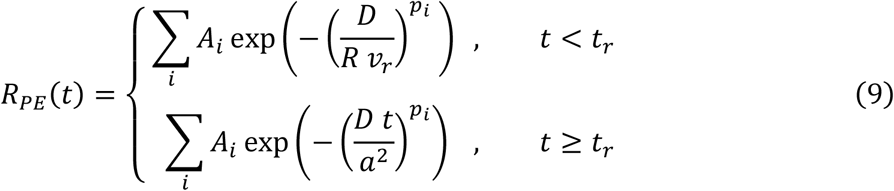

Thus, during the ramp, the poroelastic relaxation function gives a constant value which magnifies the elastic load. The level of magnification depends on the velocity of loading. This form is valid for different representations of poroelastic relaxation. It should also be noted that the poroelastic relaxation function is dependent on the final depth of indentation, via *v*_*r*_.

#### 2.1.2 Poroviscoelastic Theoretical Framework

As noted in the introduction, the solid network of soft hydrated materials may also exhibit time dependent behavior independent of the interstitial fluid flow. This intrinsic viscoelastic behavior is not reflected in the poroelastic load relaxation equation above, and can occur on the same time scale as the poroelastic relaxation. To account for this behavior of the solid matrix, we formulate a poroviscoelastic theoretical framework that builds on the Wang poroelastic load relaxation equations using the elastic-viscoelastic correspondence principle, which may be applied when the contact area monotonically increases or stays constant with time such as ramp-and-hold load relaxation [38]. First, we re-write the poroelastic force expression as follows:

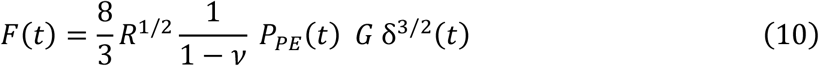

Separating this expression into ramp and hold phases of indentation and substituting *δ*(*t*) = *v*_*r*_ *t* gives the following:

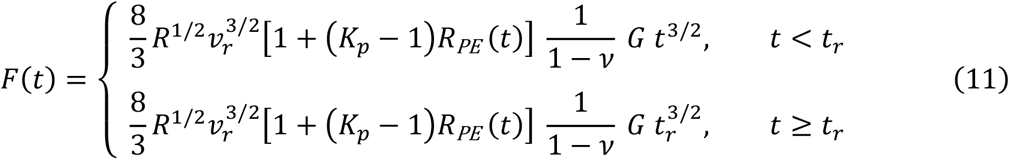

With a viscoelastic material, the shear modulus of the solid matrix is a function of time instead of a constant value. To obtain the force under load relaxation, we employ the elastic-viscoelastic correspondence principle by replacing instances of multiplication of the now transient material coefficients (shear modulus) with a function of time that appear in the poroelastic force equation with a Steiltjes convolution product [39].

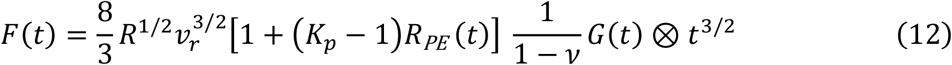

The following expression captures the poroviscoelastic force in integral form and separated into the ramp and hold phases of indentation:

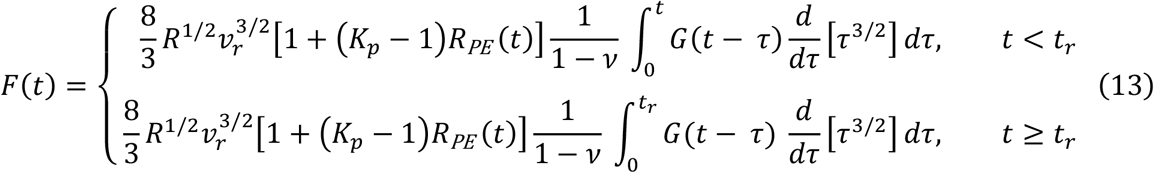

*τ* being dummy variable introduced by the convolution. We assume that the shear modulus function G(t) takes the following form:

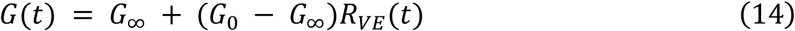

Where *G*_∞_ is the equilibrium shear modulus, *G*_0_ is the initial shear modulus, and *R*_*VE*_(*t*) is the viscoelastic relaxation function that ranges from a value of 1 at t = 0 and 0 as t approaches infinity. This represents the indenter being held at a constant displacement. *G*_∞_ and *G*_0_ are intrinsic material properties of the solid matrix. Substituting Eq 14 into Eq 13 yields the following:

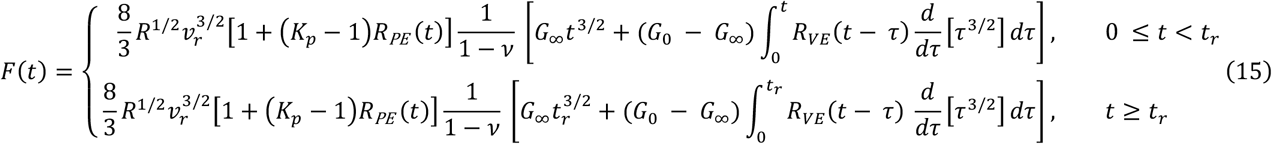

We can see in the prior equation that, if the solid matrix is elastic, *G*_0_ = *G*_∞_ and we recover the poroelastic formulation. Similarly, if there is no poroelastic contribution to relaxation (the pressurization factor *K*_*p*_ = 1, this equation reduces to a purely viscoelastic form.

#### 2.1.3 Theoretical Basis for Sequential Indentation to Isolate Poroelastic and Viscoelastic Mechanisms of Load Relaxation

Microscale indentation permits the rapid evaluation of time-dependent mechanical responses as load relaxation at this scale for soft hydrated materials approaches equilibrium on the order of seconds [34]. While this rapid experimental timeframe presents practical utility for characterization, poroelastic and viscoelastic responses that occur on the same time scale cannot be distinguished with a single test. However, we can use sequential indentation tests at the same scale but different depths to isolate poroelastic and viscoelastic mechanisms of relaxation by taking advantage of the length-scale dependence of poroelasticity and the length scale independence of viscoelasticity.

For the sequential indentation scenario, we assume an initial ramp indentation and hold at depth *δ*_1_ and a second ramp indentation and hold at different depth *δ*_2_ following sufficient recovery time. Both indentations occur at the same speed *v*_*r*_. We assume that the material constants do not vary with indentation depth. Focusing on the relaxation portion of the load curve, for each indentation represented by subscript *j*, we can generate an expression that describes normalized poroviscoelastic relaxation.

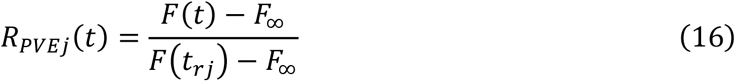

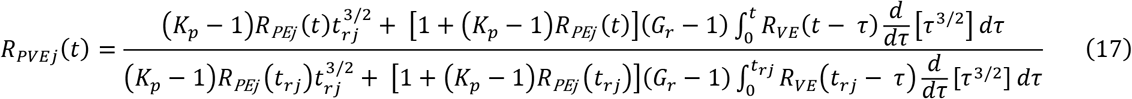

Where viscoelastic ratio *G*_*r*_ = *G*_0_/*G*_∞_. In addition to these normalized relaxation equations, we develop two additional equations that must be satisfied to extract material parameters by normalizing the maximum load which occurs at *F*_*j*_(*t*_*rj*_) with the equilibrium forces *F*_∞*j*_.

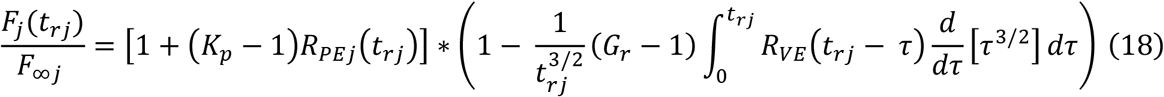

Note that *R*_*VE*_ (*t*) is the same for both indentation depths, while *R*_*PEj*_ (*t*) differs. These equations in addition to the load equations provide the necessary information to extract the poroelastic and viscoelastic properties of the tested material. While these equations cannot be solved analytically, the force, the indentation, the indenter radius, and the indentation rate are known experimentally. Thus, numerical procedures can be used to obtain unknown material constants if when the appropriate constitutive models of poroelastic and viscoelastic relaxation are provided.

#### 2.1.4 Viscoelastic Relaxation Constitutive Model

In this work, the viscoelastic properties of soft biological tissues were approximated using a generalized standard linear solid model, which captures viscoelastic relaxation behavior through an expanded sum of exponentials [34,40]. Thus, the viscoelastic relaxation function for this work is given by:

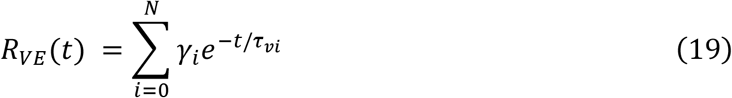

where *γ*_*i*_ is the fraction of the total stiffness dissipated (equal to *G*_0_ − *G*_∞_) at a characteristic relaxation time scale *τ*_*vi*_. This form of the relaxation function requires that ∑*γ*_*i*_ = 1. Upon substituting this constitutive form of the viscoelastic relaxation function into the poroelastic force equation (15):

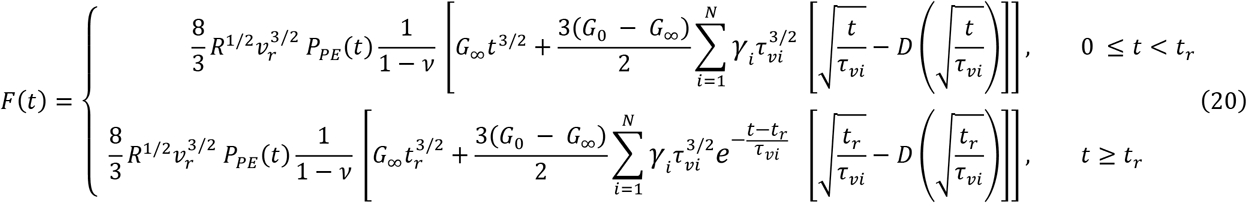

where *D*(*x*) is Dawson’s function defined as:

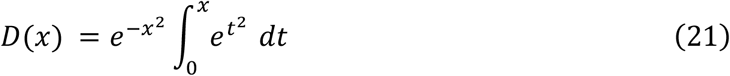

### 2.2 Theoretical Work: Completing Our Semi-analytical PVE Model and Assessing its Performance

#### 2.2.1 Empirical Derivation of Poroelastic Relaxation Model

Expanding on theoretical foundations presented by Hui et al [41] for plane-strain indentation using a plane-strain cylindrical indenter, Hu et al derived a set of empirical equations for normalized poroelastic force relaxation for a variety of indenter shapes (cylindrical, conical and spherical) from finite element simulations using ABAQUS [42]. Each of these equations is a function of normalized time (*τ* = *D t*/*a*^2^) containing two exponential terms. Later work by Lai and Hu [43] demonstrates that the normalized poroelastic relaxation for cylindrical punch indentation is better captured using an expanded four-term fitting equation. We use a similar approach to a derive a four-term fitting equation for spherical indentation to model poroelastic relaxation in this study. We created quasi-axisymmetric models of spherical indentation in FEBio (version 2.6.0) using the Biphasic Physics module with the same geometric parameters using in [42]. A soft hydrated material sample was modeled as a 3° wedge of an idealized cylindrical geometry with a 225 µm radius and 225 µm height. The spherical indenter was modeled as a one octant of a hollow spherical rigid body with a 50 µm outer radius. The soft hydrated material geometry was meshed with linear hexahedral elements using a substantial bias in which the region near the indentation contact site was more refined. A visualization is provided in Figure 1.

**Figure 1.**
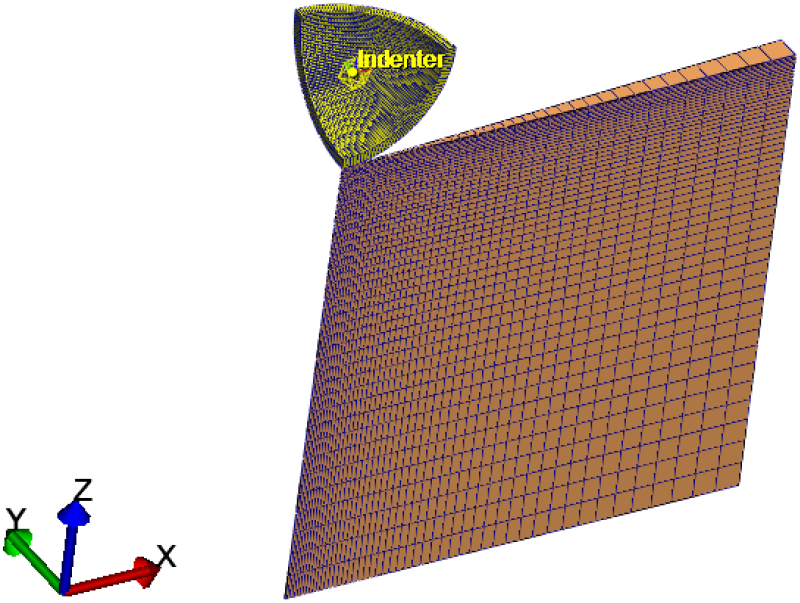
Baseline FeBio Model used for empirical poroelastic relation calculation and benchmark studies

We defined the boundary conditions by fixing all displacements of the sample’s bottom surface, fixing x- and y-displacements of the center of the sample, and applying symmetry planes to the sectioned faces of the wedge with a penalty value of 10^−6^. The outer edge of the wedge was left unconstrained. The sliding-biphasic contact model was used to describe the contact between the rigid indenter and the sample with auto penalty and augmented Lagrangian enforcement selected. The sample surface was assigned to be the primary contact surface and the indenter’s contact surface was assigned as the secondary. The external surface of the wedge was prescribed a zero-pressure boundary and initial condition to permit permeability that the surface. A Neo-Hookean material model was used for the solid phase. A mesh refinement study was conducted to determine the appropriate mesh size and resulted in 50 elements in the vertical and radial directions. The indenter motion was restricted to indentation in the z-direction to a prescribed displacement of 10 µm. The model was run four times varying the permeability by decade values (k = 0.0001 to 0.1 µm^4^/nN s) up to a normalized time of *τ* = 10^6^. These permeability values were selected to ensure that the poroelastic time constant would be much greater than the ramp time, thus reducing the effect of the initial loading rate. The reaction force in the Z-direction acting on the indenter was extracted and normalized. We used a custom Python script with a differential evolution algorithm to minimize residuals and fit the following equation:

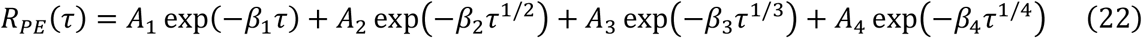

The optimization algorithm was run 5000 times with different initial seed points to find the global optimum values. These poroelastic relaxation equations were used in our subsequent semi-analytical models.

#### 2.2.2 Benchmarking Semi-Analytical PVE Model to Finite Element Models

We benchmarked the results of the semi-analytical PVE model formulation to a poroviscoelastic finite element model of soft hydrated material indentation. We used the same geometric configuration described in the prior section to create poroviscoelastic models in FEBio. Again, the Biphasic Physics module was used. In this case, the material behavior of the solid component was described by FEBio’s viscoelastic material model. We conducted the finite element simulations in two steps: an indentation ramp of 6 µm or 10 µm for 1 s followed by a hold period of 30 s. Both steps were transient simulations performed using the Full Newton solver. The material parameters used for the benchmark study were as follows: *E*_0_ = 10 *kPa, E*_∞_ = 5 *kPa*, *v*_*d*_ = 0.3, *γ*_1_ = 0.3, *γ*_2_ = 0.5, *γ*_3_ = 0.2, *k*_*h*_ = 10 *μm*^4^/*nN s*, where k is the hydraulic permeability and *v*_*d*_ is the drained Poisson’s ratio. We related the diffusivity of the semi-analytical model to hydraulic permeability in FEBio using the following equation [42,44]:

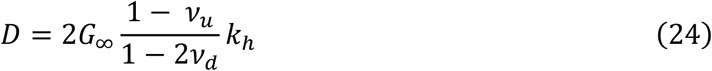

We set *v*_*u*_ = 0.5 (undrained Poisson’s ratio) as done in prior work [34,45]. We first ran the model as purely poroelastic in order to calculate *K*_*p*_. Then then ran the fully poroviscoelastic model. We compared the numerical output of the indenter reaction force in the Z-direction from the finite element simulations to the force output of the appropriate version semi-analytical PVE model. The Hertz load in the semi-analytical model was modified to account for the finite thickness using the correction formula developed by Dimitriadis et al [46]. We calculated percentage error in peak force and equilibrium force and calculated the normalized root mean square error for the entire load response.

### 2.3 Experimental Work: Application of Semi-analytical PVE Model to Soft Tissues and Hydrogels

#### 2.3.1 Experimental Samples

Fresh and healthy porcine tissue hearts and livers were obtained from a local commercial abattoir (Mary’s Ranch, Hialeah, FL). All animals were six to seven months old. The abattoir does not provide biological sex of animals and therefore unknown for these samples. Organ tissues were cut into 0.5 mm slices using a commercial meat slicer. For testing, tissue samples were glued onto dishes to ensure attached and to prevent potential slipping during loading. Tissues were submerged in phosphate buffered saline (PBS) and allowed to equilibrate at room temperature for at least 3 hours prior to testing.

Type I collagen hydrogels were formed using bovine collagen in a 3 mg/mL stock solution (PureCol ®, Advanced BioMatrix). One-part chilled neutralization buffer was added to nine parts collagen stock solution and gently mixed. The reconstituted collagen solution was then pipetted (1 mL each) into 24 well plates and placed into an incubator at 37 °C for 1 hour to allow hydrogel polymerization. Following hydrogel polymerization, collagen hydrogels were submerged in PBS overnight to equilibrate prior to testing.

Gelatin methacryloyl (GelMA) was synthesized according to previously described methods [47]. Briefly, 20 g of type A gelatin from porcine skin (300 Bloom, Sigma Aldrich) was added to 200 mL of a 0.25 M carbonate buffer solution at 50 °C for 1 hour under constant stirring. After complete dissolution of gelatin, the pH of the solution was adjusted to 9 and 0.1 mL of methacrylic anhydride was added for each gram of gelatin dropwise to the dissolved gelatin solution and allowed to react for 1 hour. Upon completion of reaction, the pH was adjusted to 7.4 and the reaction was terminated by 2x dilution with PBS. Reaction products were dialyzed against deionized water at 50 °C for three days with changes in water twice a day then lyophilized to obtain the pure gelatin methacryloyl. The dried gelatin methacryloyl was stored at -20 °C. GelMA hydrogels were prepared at 10 wt% (w/v) by dissolving GelMA in PBS at 37 °C and adding the photoinitiator lithium acylphosphinate (Chemscene) at 0.1 wt%. The solution was pipetted (1 mL) into 24 well plates and hydrogels were formed by polymerization under 405 nm light for 5 minutes. Following hydrogel polymerization, GelMA hydrogels were submerged in PBS overnight to equilibrate prior to testing.

#### 2.3.2 Sequential Microscale Relaxation Indentation Testing

Microscale indentation testing was conducted using an instrumented indentation device with spherical indentation probes (Chiaro, Optics11 Life). For all samples (n = 4 for all soft hydrated materials), a 49.5 *μm* radius probe with a cantilever stiffness of 0.42 N/m were used for testing. Following equilibration, soft hydrated material samples were indented to a fixed depth of 6 *μm* (*δ*_1_) under indentation control at a loading rate of 6 *μm*/*s*. After reaching the prescribed indentation depth, the load relaxation was performed by holding indentation for 15s. Each sample was tested using a 2 x 3 matrix scan with 250 *μm* between indentation points. Following the initial indentation scan, at the same points, samples were indented to a fixed depth of 10 *μm* (*δ*_2_) at the same loading rate of 6 *μm*/*s*.

#### 2.3.3 Determination of Microscale Time-Dependent Mechanical Properties: Analysis of Sequential Microscale Relaxation Indentation Data

Based on the derived semi-analytical poroviscoelastic model for spherical indentation, the equations that describe normalized load relaxation at each indentation depth are given Equation 17 and repeated here:

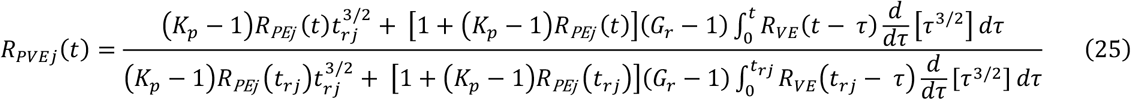

For the Prony series constitutive description of viscoelasticity, we select a series consisting of three terms. We fix the relaxation time scales *τ*_*i*_ in the Prony series a priori as done in prior work, searching only for the normalized relaxation moduli which determine the strength of each time scale. The characteristic time constants selected (*τ*_1_ = 0.15 *s, τ*_2_ = 1.5 *s*, and *τ*_3_ = 15 *s*) to cover the time scale of the experiment. The six material parameters to be identified are *K*_*p*_, *D, G*_*r*_, *γ*_1_, *γ*_2_, and *γ*_3_. These parameters were determined by curve-fitting using custom developed Python script. The script fits the normalized load relaxation curves using a differential evolution algorithm from SciPy’s optimization library [48] where *r*^2^ values of the model vs experimental data were maximized. The large parameter space was reduced through Equations 18, transforming that equation into inequality constraints as shown in Equation 26. Any parameter combination that violated this constraint was rejected regardless of fit to the load relaxation equations.

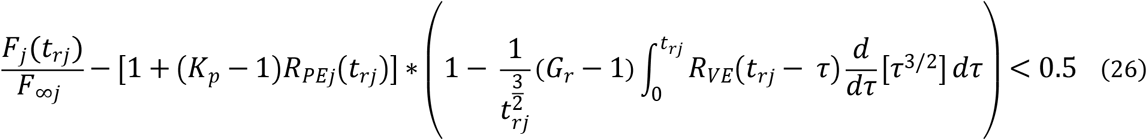

#### 2.3.4 Statistical Analysis

Poroviscoelastic parameters were analyzed for each of the soft hydrated materials tested using one-way ANOVA with the assumption of normal data distribution and homogeneity of variances. For diffusivity, a log transformation was applied to meet normality assumptions. Tukey’s Honestly Significant Difference (HSD) test was used for post-hoc comparisons where ANOVA indicated significant differences (p < 0.05). All data analysis was conducted using Python, with the Pandas library for data manipulation, the Statsmodels library for statistical testing, and the Seaborn library for data visualization.

## 3 Results

### 3.1 Empirical equations for normalized poroelastic relaxation

The global optimization procedures performed in this study produced the following modified 4-term poroelastic relaxation functions:

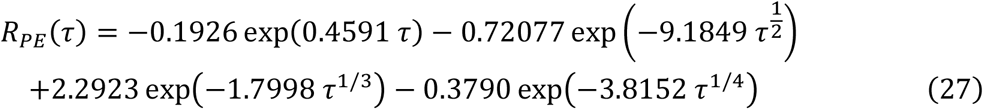

This new poroelastic relaxation functions was plotted against the FEBio simulation results over the entire normalized time scale of relaxation (from 10^−4^ to 10^6^) and over a shorter time scale representative of less than 60 s of relaxation for *k*_*h*_ = 10 *μm*^4^/*nN s*. The results are plotted in Figure 2. Over the entire time domain, the 4-term poroelastic relaxation model demonstrated great fit accuracy with a NRMSE of 0.002 for the entire time range and 0.003 for the short-term normalized time values. These values were an order of magnitude lower than the original two-term Hu equation. This difference is particularly pronounced in early normalized time ranges which is critical for nanoindentation.

**Figure 2.**
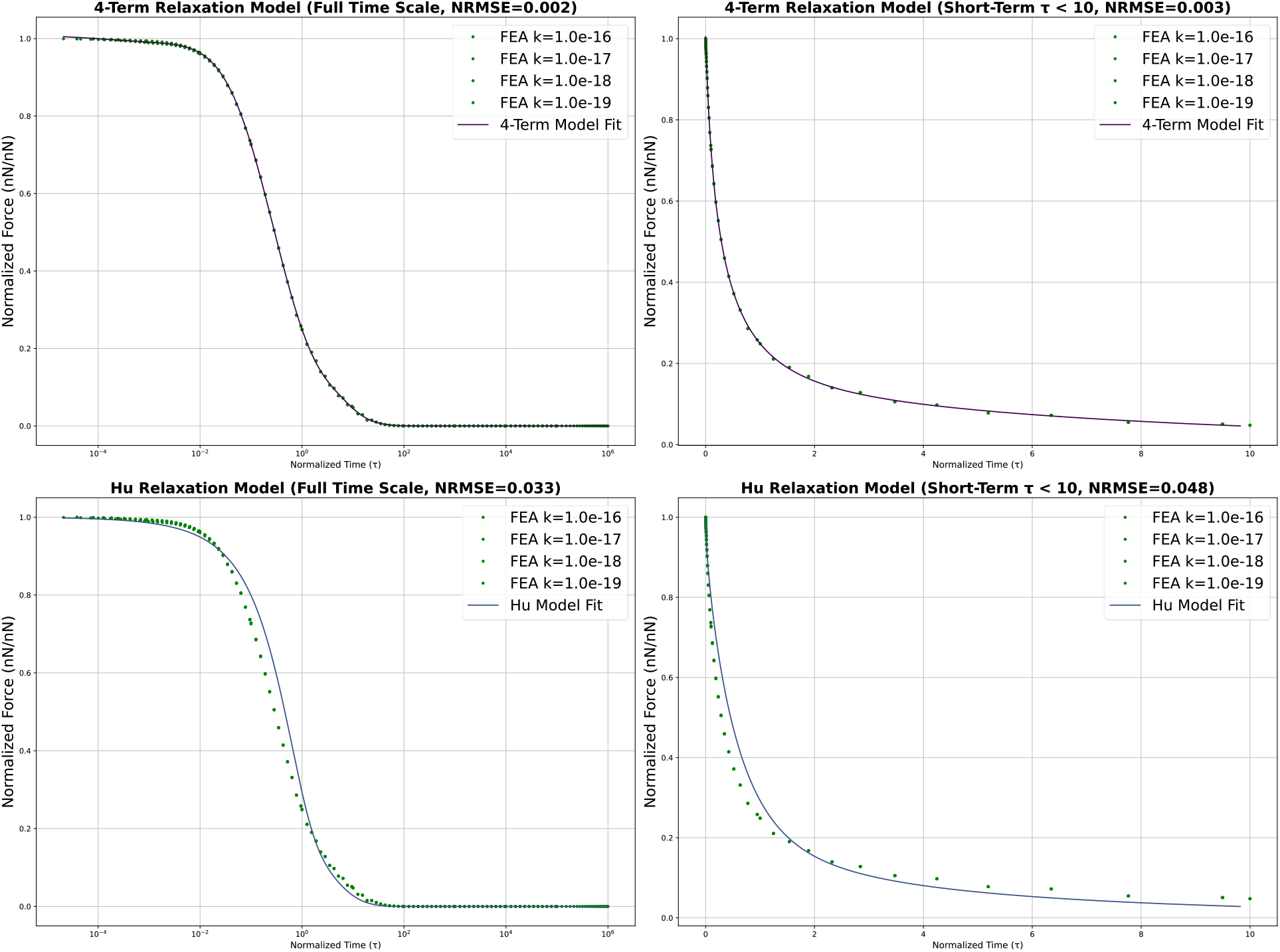
Comparison of Relaxation Models Fitted to Finite Element Analysis (FEA) Data at Different Timescales. This figure presents normalized force relaxation data fitted using two models: a 4-Term Normalized Relaxation Model and the Hu Normalized Relaxation Model. Each model is assessed over a full-time scale and a short-term scale where normalized time *τ* is less than 10. The normalized root-mean-square error (NRMSE) values are provided for each model fit, indicating the goodness of fit. The data points represent finite element analysis (FEA) results for various values of the hydraulic permeability *k* (from 1 × 10^−16^ to 1 × 10^−19^). The fitted model curves are overlaid to show the models’ accuracy in predicting the force relaxation behavior over time.

### 3.2 The semi-analytical model matches finite element simulations for poroviscoelastic relaxation

To determine if our semi-analytical model could effectively represent soft hydrated material behavior with overlapping poroelastic and viscoelastic relaxation, we compared the model results with load relaxation curves from finite element models of spherical indentation. The semi-analytical PVE model fit the finite element model force output very well for both indentation depths (*δ*_1_ = 6 *μm* and *δ*_2_ = 10 *μm*) and for both the indentation ramp and isometric hold periods (Figure 3). The normalized root mean square error (NRMSE) of the indentation ramp for the 6 *μm* and the 10 *μm* simulations were 0.273 *x* 10^−2^ and 0.725 *x* 10^−2^, respectively. Similar low NRMSE values were measured for the isometric hold portion of the curve (1.564 *x* 10^−2^ and 0.810 *x* 10^−2^, respectively).

**Figure 3.**
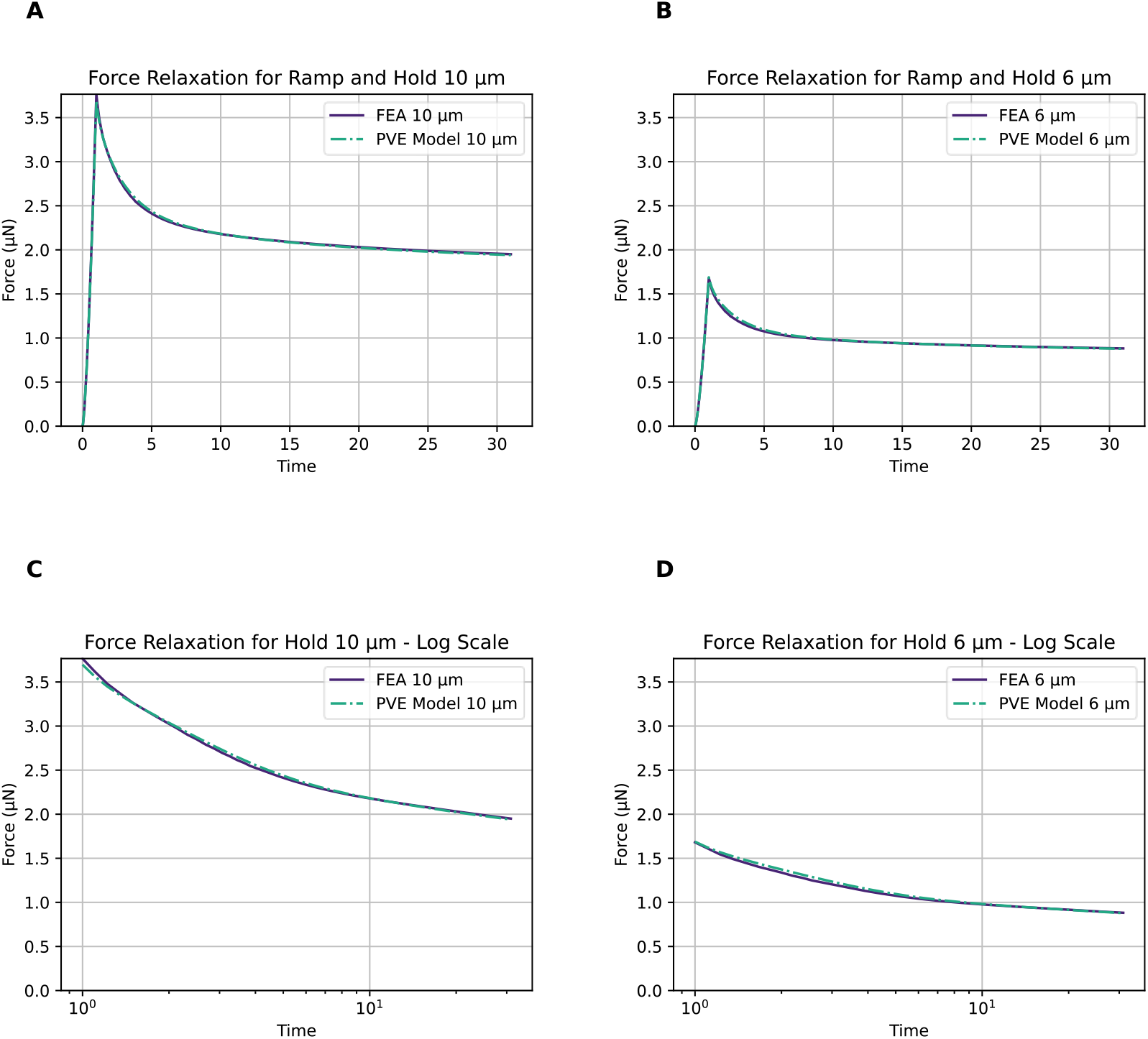
Benchmark analysis of semi-analytical PVE model: Force relaxation curves for indentation ramp and isometric hold experiments at indents of 10 μm and 6 μm simulated by finite element analysis and compared to the semi-analytical results. All models show the expected initial rapid drop in load followed by a more graduate relaxation to equilibrium. PVE model shows excellent agreement with FEA simulations in capturing poroviscoelastic behavior of soft hydrated materials at different indentation depths.

### 3.3 Semi-analytical PVE model response for different material types

We investigated how the semi-analytical poroviscoelastic model proposed in this paper described the load response during spherical indentation consisting of a ramp followed by an isometric hold. By manipulating the pressurization constant (*K*_*p*_) and the viscoelastic ratio (Gr), we explored four different material types: 1) elastic material *K*_*p*_ = 1, *G*_*r*_ = 1), 2) poroelastic material (*K*_*p*_ > 1, *G*_*r*_ = 1), 3) viscoelastic material (*K*_*p*_ = 1, *G*_*r*_ > 1), and 4) poroviscoelastic material (*K*_*p*_ > 1, *G*_*r*_ > 1). We also varied the ramp time (*t*_*r*_) by decade values while maintaining the same indentation depth to explore how the model described the behavior of the different materials under varying indentation rates (Figure 4).

**Figure 4.**
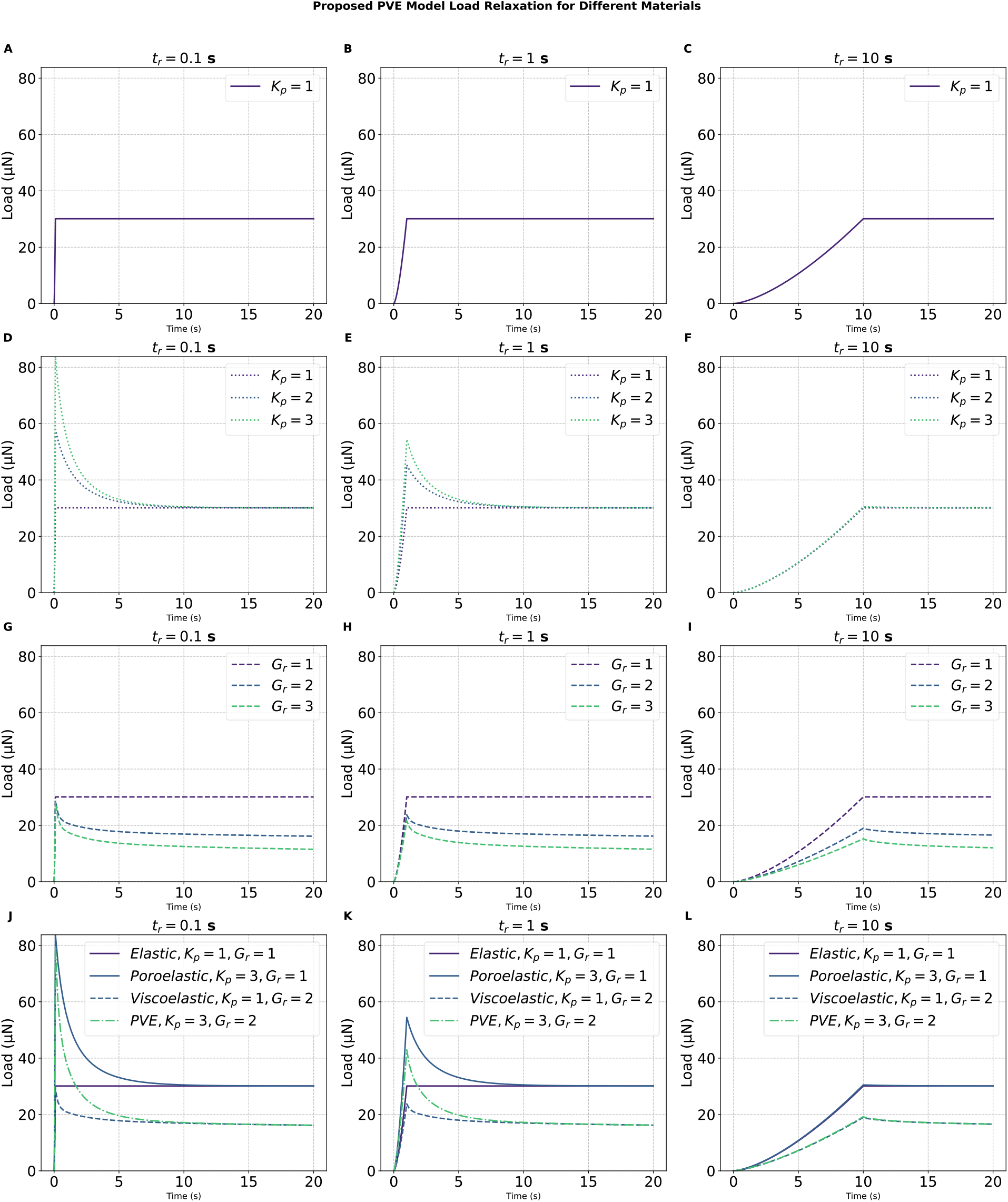
Load Relaxation Behavior of Different Materials Using the PVE Model Under Various Time-Dependent Loading Conditions. Panels (A), (B), and (C) depict the load relaxation curves for materials modeled as purely elastic (with *K*_*p*_ = 1) over three different ramp times (*t*_*r*_ = 0.1 *s*, 1 *s*, and 10 *s* respectively). Panels (D), (E), and (F) show the effect of varying the poroelastic parameter (Kp) on the load relaxation behavior for a poroelastic material with a ramp time of 0.1 *s*, 1 *s*, and 10 *s* respectively. Panels (G), (H), and (I) illustrate the impact of different viscoelastic parameters (Gr) on the load relaxation behavior for a viscoelastic material across the same ramp times. Panels (J), (K), and (L) compare the load relaxation behavior of elastic, poroelastic, and viscoelastic materials, as well as a combined PVE material with different parameter settings, highlighting the complex interactions and relaxation characteristics under varying ramp times of 0.1 *s*, 1 *s*, and 10 *s*.

In the case of an elastic material, the load vs time response for the ramp is equivalent to the Hertz formulation for indentation. Further, during the isometric hold portion of the curve, the load remains constant. This behavior is depicted in Figure 4 A – C, showing a consistent load level across varying time ramps (Figure 4A – C). In poroelastic materials, the model reflected intuitive load relaxation behavior, with a higher peak loading for higher pressurization constant. Additionally, as illustrated in Figure 4 D – F shorter ramp times (higher indentation rates) led to higher peak loads. Similarly, for purely viscoelastic materials, the model predicts that a shorter ramp time (higher indentation rate) results in a higher peak load. However, unlike poroelastic materials, a purely viscoelastic material does not experience a pressurization over the Hertz load. Instead, for materials with the same initial shear modulus, the peak load approaches but does not quite reach the “ceiling” Hertz value given by the initial shear modulus *G*_0_. As shown in Figure 4 G – I, as time progresses, under isometric hold, load relaxation proceeds down to the Hertz load value given by the equilibrium shear modulus *G*_∞_ . Further, with a longer ramp time, the more dissipation occurs during the ramp. The model demonstrates that poroviscoelastic materials exhibit an initial pressurization near the poroelastic peak load (for the same value of *K*_*p*_) and relax down to the viscoelastic equilibrium load. This combined behavior is shown in Figure 4 J – L, highlighting the model’s capability to represent both fluid flow and viscoelastic dissipation.

### 3.4 Experimental Results: Identification of Poroelastic and Viscoelastic Mechanical Properties through Sequential Indentation

In this study, we used sequential indentation to characterize the poroelastic and viscoelastic material properties of two commonly used biomaterial hydrogels (GelMA and collagen) and two soft biological tissues (porcine heart and liver). Representative graphs of sequential indentation are shown in Figure 5. The curves are fully normalized using Equation 17. Curve fits using the PVE model for both curves are also shown in Figure 5. All samples underwent rapid relaxation within the short time interval with the exception of GelMA samples which required more time to reach equilibrium (30s).

**Figure 5.**
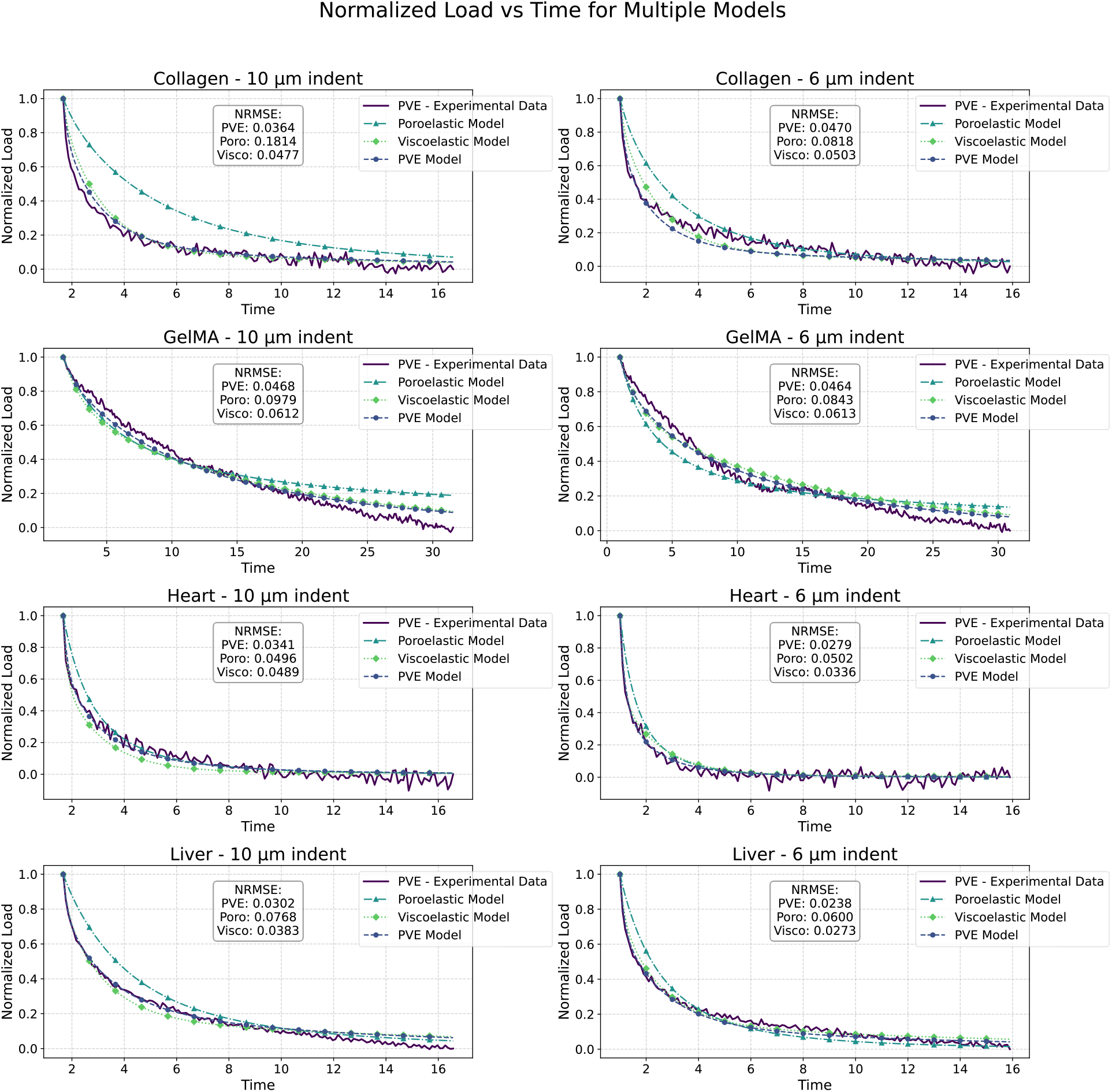
Illustrations of the normalized load as a function of time for four different soft hydrated materials: Collagen, GelMA, Heart, and Liver, each subjected to two indentation sizes, 10 μm (left column) and 6 μm (right column). The plots compare experimental data with three different model predictions: PVE (poroviscoelastic), poroelastic, and viscoelastic. Experimental data, denoted as PVE - Experimental Data, are shown with solid lines. PVE Model predictions are depicted with dashed lines and circle markers, Poroelastic Model predictions with dash-dot lines and triangle markers, and Viscoelastic Model predictions with dotted lines and diamond markers. The Normalized Root Mean Square Error (NRMSE) for each model is annotated on the plots, providing a measure of model accuracy relative to the experimental data. Lower NRMSE values indicate better model performance in replicating the observed data.

We used custom optimization software to perform parameter identification of the parameters by fitting the normalized isometric hold regions of the sequential indents of soft hydrated materials to the semi-analytical PVE model. The custom optimization software repeated the search for 50 iterations to identify the parameters which best fit the model. The poroelastic parameters included the pressurization constant (*K*_*p*_) and the diffusivity (*D*). The viscoelastic parameters extracted were the viscoelastic ratio (*G*_*r*_ = *G*_0_/*G*_∞_), and the normalized viscoelastic relaxation moduli (*γ*_1_, *γ*_2_, and *γ*_3_). In Table 1, we present the poroelastic and viscoelastic parameters extracted from the sequential indentation tests for each of the soft hydrated tissues. Additionally, Figure 6 shows a comparative analysis of the poroelastic and viscoelastic properties across the different soft hydrated materials in this study.

**Table 1:**
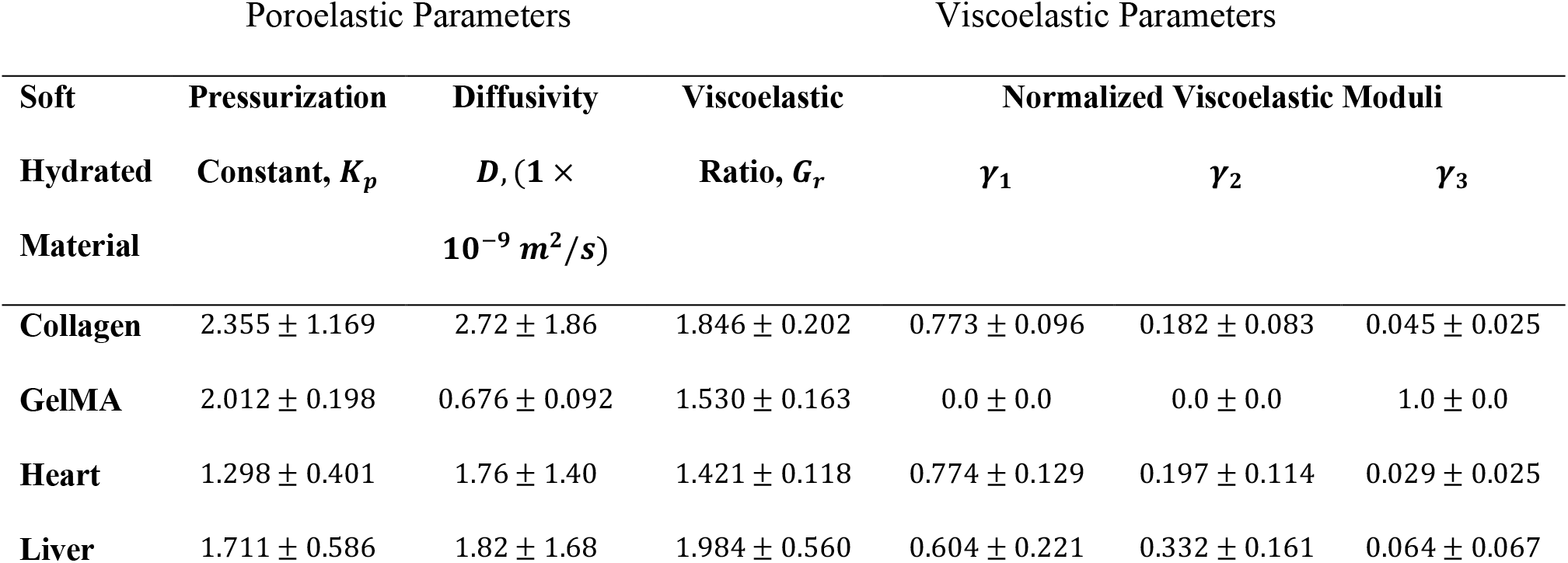
Poroelastic and Viscoelastic Parameters of Soft Hydrated Materials as Determined by Sequential Nanoindentation (Mean +/-St. Dev.)

| Poroelastic Parameters |  |  | Viscoelastic Parameters |  |  |  |
| --- | --- | --- | --- | --- | --- | --- |
| Soft | Pressurization | Diffusivity | Viscoelastic | Normalized Viscoelastic Moduli |  |  |
| Hydrated | Constant, $K_p$ | $D$, ($1 \times$ | Ratio, $G_r$ | $\gamma_1$ | $\gamma_2$ | $\gamma_3$ |
| Material | | $10^{-9} m^2/s$) | | | | |
| Collagen | $2.355 \pm 1.169$ | $2.72 \pm 1.86$ | $1.846 \pm 0.202$ | $0.773 \pm 0.096$ | $0.182 \pm 0.083$ | $0.045 \pm 0.025$ |
| GelMA | $2.012 \pm 0.198$ | $0.676 \pm 0.092$ | $1.530 \pm 0.163$ | $0.0 \pm 0.0$ | $0.0 \pm 0.0$ | $1.0 \pm 0.0$ |
| Heart | $1.298 \pm 0.401$ | $1.76 \pm 1.40$ | $1.421 \pm 0.118$ | $0.774 \pm 0.129$ | $0.197 \pm 0.114$ | $0.029 \pm 0.025$ |
| Liver | $1.711 \pm 0.586$ | $1.82 \pm 1.68$ | $1.984 \pm 0.560$ | $0.604 \pm 0.221$ | $0.332 \pm 0.161$ | $0.064 \pm 0.067$ |

**Figure 6.**
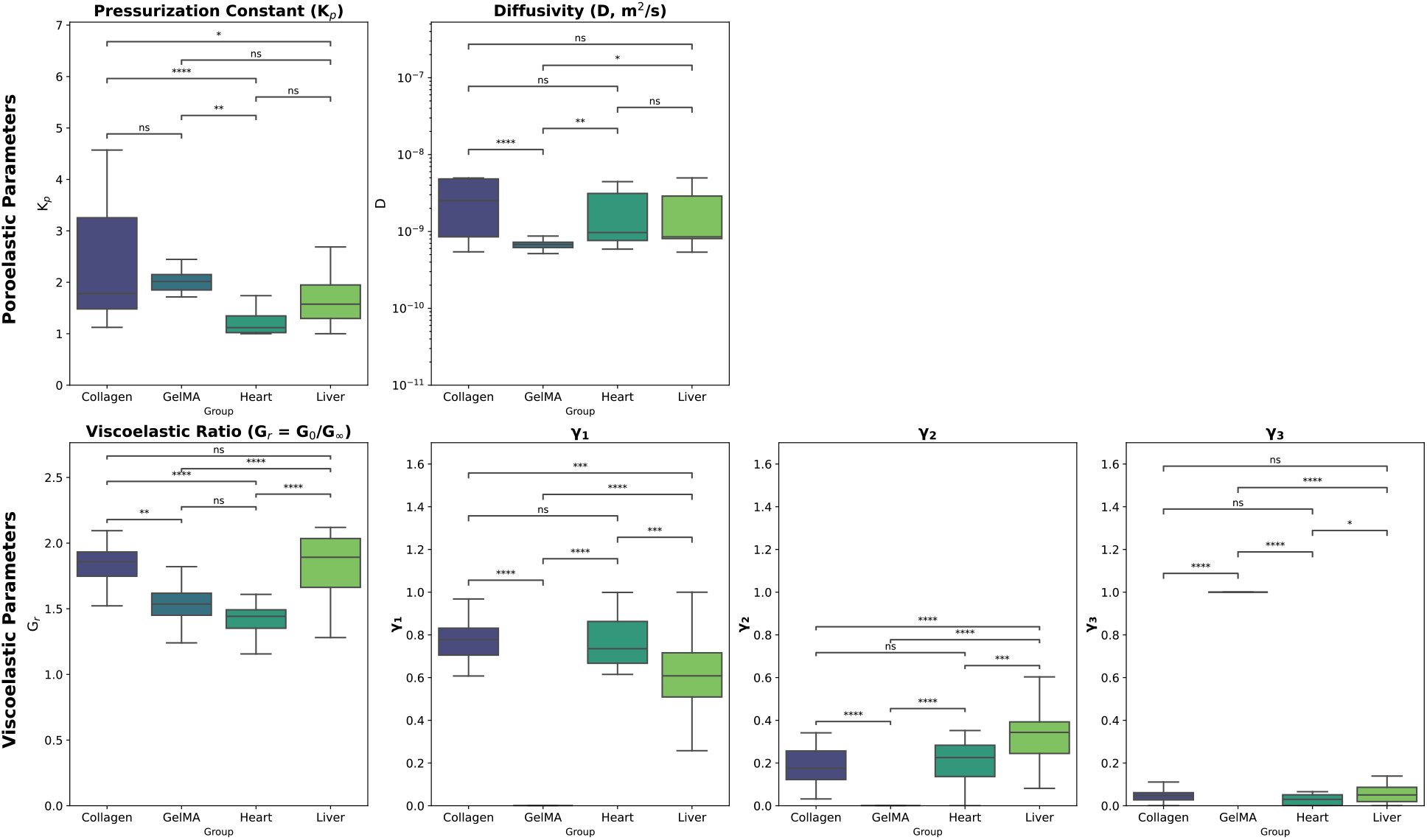
Box plots depicting the distribution of various poroelastic and viscoelastic parameters across four different soft hydrated materials: Collagen, GelMA, Heart, and Liver. The parameters evaluated are *K*_*p*_ (pressurization constant), *D* (diffusivity), *G*_*r*_ (viscoelastic ratio), *γ*_1_, *γ*_2_, and *γ*_3_ (normalized viscoelastic moduli). Each plot illustrates the median, interquartile range, and outliers for the respective parameters across the material groups. The statistical significance of pairwise comparisons is annotated, where ns denotes non-significant, * *p* < 0.05, ** *p* < 0.01, *** *p* < 0.001, and **** *p* < 0.0001. The y-axes for *γ*_1_, *γ*_2_, and *γ*_3_ are set to a common scale to facilitate direct comparison.

Comparative analysis of the different soft hydrated materials characterized through sequential indentation in this study reveals some significant differences in the poroelastic and viscoelastic behaviors. First, while collagen and GelMA hydrogels possess similar pressurization constant values, collagen has a *K*_*p*_ value (2.355 ± 1.169) that is significantly higher than the soft tissues tested, indicating a higher load amplification and a greater magnitude of poroelastic relaxation. GelMA’s *K*_*p*_ value (2.355 ± 1.169) is significantly higher than heart (*K*_*p*_ = 2.012 ± 0.198) but not statistically different from liver (*K*_*p*_ = 1.711 ± 0.586). The diffusivity values, however, varied between the two hydrogels. Our results show that collagen has an order of magnitude higher diffusivity than GelMA. GelMA’s diffusivity was also significantly lower than that of heart and liver tissue. The viscoelastic behavior varied notably between the soft hydrated materials. First, in examining the viscoelastic ratio, GelMA (*G*_*r*_ = 1.530 ± 0.163) and heart tissue (*G*_*r*_ = 1.420 ± 0.118) exhibited a lower viscoelastic ratio, indicating more elastic behavior than collagen (*G*_*r*_ = 1.846 ± 0.201) and liver (*G*_*r*_ = 1.984 ± 0.560). The normalized viscoelastic relaxation moduli (*γ*_1_, *γ*_2_, and *γ*_3_) represent the relative time scales of viscoelastic relaxation in the materials. The viscoelastic relaxation moduli of the earlier time scales (*γ*_1_ and *γ*_2_) were notably found to be 0 for all GelMA samples. This is substantially different from the other three materials. Collagen and heart have rather high early normalized viscoelastic relaxation moduli (*γ*_1_ ≈ 0.78) and similar mid-range normalized relaxation moduli (*γ*_2_ ≈ 0.18). The three liver moduli are more equally weighted between the two early time scales.

Lastly, we also examined the PVE model fit for sequential indents compared to poroelastic and viscoelastic model fits to the same indentation data using the poroelastic and viscoelastic constitutive equations that are used in our model. Figure 5 shows the average fit for the different models. Figure 7 provides the distribution of *r*^2^ values for each of the two indentation curves using the three models. The results indicate that, for all soft hydrated materials, that the PVE model produces a better fit to the relaxation data (Figure 7).

**Figure 7.**
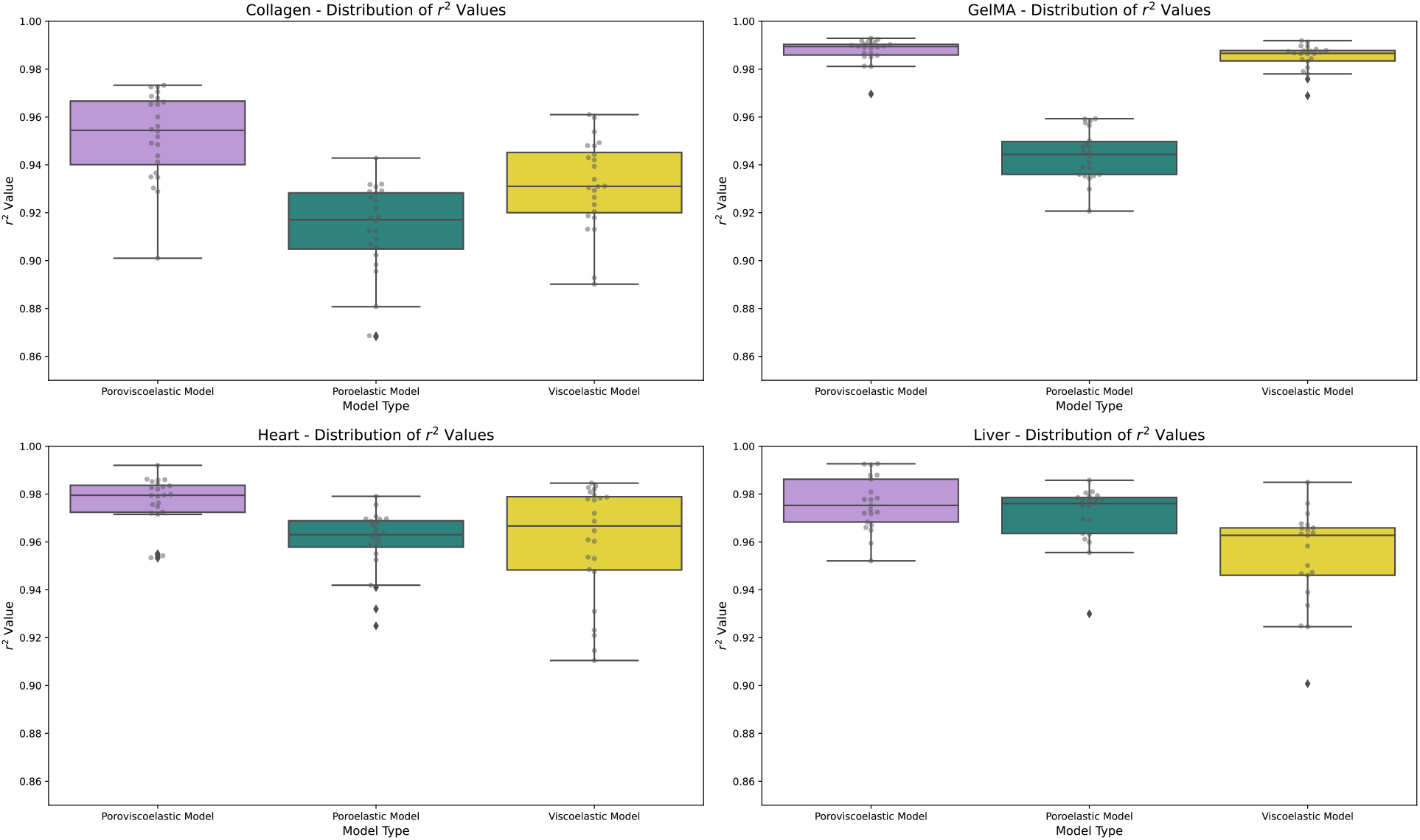
Distribution of r^2^ values for Different Models Across Various Materials. This figure displays the distribution of *r*^2^ values for three different models—Poroviscoelastic, Poroelastic, and Viscoelastic—across four materials: Collagen, GelMA, Heart, and Liver. Each subplot represents a specific material and shows the range and median of *r*^2^ values obtained from fitting the respective model to the data. The *r*^2^ values are presented on a scale from 0.85 to 1.0, indicating the goodness of fit of the hold region for both the 6 *μm* and the 10 *μm* indents in the sequential indentation procedure. Higher *r*^2^ values denoting better model performance. The light purple, teal, and yellow box plots correspond to the Poroviscoelastic, Poroelastic, and Viscoelastic models, respectively. The data points are overlaid as dots for clarity. The box plots highlight the variability and central tendency of the *r*^2^ values for each model and material combination, providing a comparative view of model performance across different materials.

## 4 Discussion

In this study, we presented a new semi-analytical poroviscoelastic (PVE) model for characterizing the time-dependent mechanical behaviors of soft hydrated materials under spherical indentation. Our work extends existing poroelastic models and offers a rapid and robust approach to isolate poroelastic and viscoelastic contributions to load relaxation occurring at the same time scale. This type of characterization is critical to understanding complex material responses that occur on the size-scale of cells. From this model, we developed a mathematical formulation to use sequential microscale indentation in order to isolate poroelastic and viscoelastic relaxation properties occurring on the same time scale. We benchmarked the PVE model performance to finite element models with the same poroelastic and viscoelastic parameters. We then performed sequential nanoindentation and applied our analytical framework to quantify the poroviscoelastic properties of several different soft hydrated materials, accurately describing the load relaxation response for both indentation levels occurring within 30 seconds to equilibrium. This method is particularly relevant for advancing our understanding of how cells experience these time-dependent cues within normal tissues and within simulated environments, as highlighted in the introduction.

The formulation of our semi-analytical PVE model provides a conceptual framework to understand poroviscoelastic behavior of soft hydrated materials. First, in the process of deriving the model, we implemented a new mathematical representation for load amplification in poroelasticity captured by the pressurization constant term (*K*_*p*_). Mathematical analysis of the PVE model reveals that *K*_*p*_ is a multiplier for elastic load value produced by a hypothetically applied step displacement. This amplified load (*K*_*p*_ *F*_*elastic*_) represents the maximum load amplification that can occur. Given that a true step indentation cannot be applied, the actual pressurization value will be less than *K*_*p*_ due to the outgoing flow of fluid during the ramp indentation. While one can choose to ignore the fluid flow, when diffusivity is in the range commonly reported for hydrogels and soft tissues (on the order of 10^−7^ − 10^−12^ *m*^2^/*s*), significant fluid flow can occur in the ramp time of just 1 second [37,44,49,50]. We find that the amount of dampening to poroelastic pressurization is related to the indenter radius, diffusivity, and the velocity of the ramp. Also notable in the formulation is the constant value of pressurization during the ramp loading. During the isometric hold stage, the pressurized fluid will diffuse away from the indentation site leading to an equilibrium response equal to the elastic Hertz load.

By modifying the key poroelastic and viscoelastic parameters, we demonstrated that semi-analytical PVE model effectively captures the complex load response during indentation with distinct behaviors for elastic, poroelastic, viscoelastic, and poroviscoelastic materials. The model demonstrates the poroelastic and viscoelastic material behaviors during ramp indentation and isometric hold are essentially the bounds of the poroviscoelastic material, with an initial pressurization near the poroelastic ceiling load and an equilibration down to the viscoelastic floor load (Figure 4 J – L). The final semi-analytical PVE model presented in this work bears some similarities to the multiplicative representation of poroviscoelasticity by Strange, Oyen, and colleagues, which describes poroviscoelastic relaxation as the multiplication of the poroelastic load times the viscoelastic load and normalized to equilibrium load [51]. Key differences lie in the representation of poroelasticity as described above and the description of both the ramp as well as hold portions of loading. These key differences may have allowed the PVE model developed in this study to have excellent agreement to experimental outputs, in addition to the benchmark FE output.

In the course deriving this PVE model in performing the initial benchmarking work, we found a mismatch in the poroelastic relaxation predicted by our model when using the Hu formula and the FEBio finite element simulation of a biphasic material. This mismatched provided the initial motivation for exploring a new formula for poroelastic relaxation. Using methods similar to those outlined in [42], we developed a new empirically derived formula for normalized poroelastic relaxation during the isometric hold – a modified 4-term relaxation function that takes the form introduced by Lai and Hu [43]. We selected this 4-term normalized poroelastic relaxation function for further use. This function incorporates fractional components the enable more accurate fitting to the finite element simulations of biphasic relaxation, particularly in the early normalized time ranges. While the changes in this time range appear small compared to later times, the subtle differences produce orders of magnitude differences in errors. Given the rapid load relaxation rate in microindentation tests, the mathematical representations of relaxation must accurately capture the expected poroelastic relaxation behavior at the early time scale.

Our study demonstrates that the semi-analytical poroviscoelastic (PVE) model is a robust tool for simulating the complex mechanical behavior of soft hydrated materials under indentation. Our benchmark analyses comparing PVE model performance in predictive ramp and loading response to FEBio simulations of spherical indentation demonstrate equivalence between the derived semi-analytical model and finite element simulations for poroviscoelastic relaxation. The force outputs from the semi-analytical PVE model closely matched those from the FE simulations, as indicated by the low normalized root mean square error (NRMSE) values. Using the constitutive relations reported above, the analytical process described for sequential indentation identified the poroelastic and viscoelastic parameters of interest through 50 optimization cycles on a standard desktop PC within one minute for each pair of sequential indents in comparison to 5 minutes for a single finite element simulation. Thus, this PVE model formulation reduces need for computationally and time intensive inverse finite element analysis to separate poroelastic and viscoelastic properties of materials.

Using the sequential indentation approach and subsequent analysis, we were able to identify poroelastic and viscoelastic parameters for four very different soft hydrated materials, including collagen and GelMA hydrogels which are commonly used as biomaterials for in vitro modeling and tissue engineering. Sequential indentation at two depths allowed us to determine that a singular mechanism of load relaxation is not sufficient to capture the observed time-dependent mechanical behavior of the soft hydrated tissues tested in this study. While the duality of relaxation mechanisms is generally acknowledged in performing time dependent mechanical characterization, this may not be evident with a single indentation, where both poroelastic and viscoelastic models can produce excellent fits to the data [34]. Interestingly, this observation holds true even for the GelMA hydrogels, where it is expected that the primarily covalent crosslinking of the structure would produce a poroelastic response. While this is the case at the earlier time periods, our model demonstrates that, at the longer time scales, viscous dissipation is required as part of the model in order to fit the indentation data. We can conclude that, at the microscale at timeframes longer than 15s, viscoelastic relaxation plays a role in stress decay. We note that in this work, we use a viscoelastic constitutive model with discrete time constants and make the common assumption that a single diffusivity is adequate for describing poroelastic response. Given the heterogeneity of biological materials, models with more complexity and a continuous relaxation spectrum may be warranted. The PVE formulation as outlined in the Theory section of this paper permits adding this kind of complexity to the model to capture a wider range of material behaviors.

The sequential indentation results also provide some insight into the structure of the materials. For example, the contrast in diffusivity between the materials can be explained by differences in pore size and structure. In comparing the two hydrogels, collagen’s higher diffusivity may result from a more open network architecture, allowing for faster fluid transport, while GelMA’s denser network restricts diffusion, leading to slower fluid movement. Such differences have been shown to impact cell viability within these constructs [52]. Additionally, it is of note that, for all parameters, GelMA hydrogels have the least variability, indicating less heterogeneity within this material compared to collagen hydrogels and the more complex biological tissues. The major differences in normalized viscoelastic moduli among the materials suggest varying degrees of interaction between their polymeric chains and the surrounding matrix. The notable absence of early-time scale viscoelastic relaxation in GelMA indicates a lack of significant short-term chain mobility, which matches our physical knowledge of GelMA structure due to its tighter cross-linking and shorter polymeric chains. The distinct poroviscoelastic properties of the hydrogels may highlight their suitability for use in different biomedical applications, particularly when comparing to natural tissues.

## 5 Conclusions

In this work, we derived a semi-analytical poroviscoelastic model to describe the time-dependent mechanical response of soft hydrated materials undergoing spherical indentation as well as an experimental approach and analytical framework to characterize both poroelastic and viscoelastic properties of the soft hydrated materials. The semi-analytical PVE model proposed, and the sequential indentation approach presented have broad application for rapid fundamental characterization of time-dependent properties of materials and understanding the underlying mechanisms that give rise to those properties. We demonstrated the utility of the model in comparison to finite element simulations. Additionally, we were able to characterize a broad range of soft hydrated materials isolating poroelastic and viscoelastic influences on relaxation using the experimental approach described. We note that the PVE model presented may be adapted by selecting different constitutive models for poroelastic and viscoelastic material behavior, which may describe even more complex materials. However, the experimental and analytical approach would remain the same. Using sequential indentation, both poroelastic and viscoelastic parameters can be rapidly and efficiently determined, which may facilitate more accurate material characterization and may also translate to measurement in clinical settings.

## CRediT authorship contribution statement

**Muhtadi Munawar Zahin**: Writing – reviewing and editing, Writing – original draft, Data curation, Formal Analysis, Methodology, Visualization

**Abeer Al Barghouthi**: Writing – reviewing and editing, Data curation, Methodology

**Darryl A. Dickerson, PhD**: Conceptualization, Writing – reviewing and editing, Writing – original draft, Data curation, Formal Analysis, Methodology, Visualization, Funding Acquisition, Project Administration, Supervision, Resources

## Data Availability

Data made available upon request.

